# H^+^- and Na^+^-elicited swift changes of the microtubule system in the biflagellated green alga *Chlamydomonas*

**DOI:** 10.1101/121137

**Authors:** Yi Liu, Mike Visetsouk, Michelle Mynlieff, Hongmin Qin, Karl F. Lechtreck, Pinfen Yang

## Abstract

The microtubule cytoskeletal system is integral to diverse cellular processes. Although microtubules are known for dynamic instability, the system is tightly controlled in typical interphase animal cells. In contrast, diverse evidence suggests that the system is mercurial in the unicellular fresh water green alga, *Chlamydomonas*, but intense autofluorescence from photosynthesis pigments has hindered the investigation. By expressing a bright fluorescent reporter protein at the endogenous level, we demonstrate in real time discreet sweeping changes in algal microtubules elicited by fluctuation of intracellular H^+^ and Na^+^. These results suggest disparate sensitivity of this vital yet delicate system in diverse organisms; and illuminate how pH may drive crucial cellular processes; how plants respond to, and perhaps sense stresses; and how many species could be susceptible to accelerated changes in global environments.

## Introduction

The microtubule (MT) cytoskeletal system is integral to many crucial processes in eukaryotic cells. The opposing ends of these cylindrical polymers exhibit distinct properties, with the plus end growing and shrinking stochastically. MTs establish the polarity of cells and serve as tracks for positioning and trafficking intracellular components. They could also form complex machineries. One example is the mitotic apparatus enabling accurate segregation of chromosomes during mitosis and meiosis. These actions require harnessing MT dynamic instability and the involvement of a wide array of accessory proteins and various post-translational modifications (reviewed by Hashimoto, 2015; Song and Brady, 2014). Not surprisingly, MTs are the target of natural toxins, such as Taxol (Weaver, 2014), and a number of herbicides (reviewed by Hashimoto, 2015). Notably, biotic and abiotic stresses alter plant MTs, while Taxol exacerbates stress-induced maldevelopment of seedlings (reviewed by Wang, et al., 2011; Hardham, 2013; Oda, 2015; Hepler, 2016). Yet the mechanisms underlying stress-induced changes of MTs and the broad implications remain elusive.

*Chlamydomonas reinhardtii* is uniquely suited for addressing these issues. The unicellular fresh water green alga has signature features of both animal and plant cells (Merchant et al., 2007). Like animal cells, it has motile flagella that contain stable MT bundles. Like typical plant cells, the alga is equipped with vacuoles, chloroplast, and dynamic cortical MTs that serve as tracks for the delivery of the enzymes which synthesize the cell wall (Paredez et al., 2006). Curiously, its MT system appears susceptible to environmental changes. Its flagella sever readily when the aqueous environment changes suddenly (Lefebvre et al., 1978; reviewed by Quarmby, 2009). The best characterized stimulus is pH shock. Organic acid, such as acetic acid (HA) diffuses across the plasma membrane and then becomes ionized. The resulting acidification in the cytosol triggers nearly simultaneous influx of extracellular Ca^2+^, which signals flagella amputation (Quarmby et al., 1996; Wheeler et al., 2008; Hilton et al., 2016). Cortical MTs become fewer and shorter, detected by immunofluorescence and biochemistry 5 mins after pH shock (Wang et al., 2013).

While live cell imaging will be ideal for revealing these acid-induced responses with higher spatial and temporal resolution, autofluorescence from photosynthetic pigments in chloroplast obscures commonly used fluorescent reporters (Lang et al., 1991; Rasala, 2013). Recently, we succeeded in revealing dynamic cortical MTs by taking advantage of the new fluorescent protein, NeonGreen (NG) that is 2.7 X brighter than EGFP (Shaner et al., 2013), and the relative abundant plus end-binding protein, EB1, as the NG carrier (Harris et al., 2016).

EB1 plays central roles in eukaryotes (Su et al., 1995; reviewed by Akhmanova and Steinmetz, 2010; Kumar and Wittmann, 2012). Its N-terminal domain preferentially binds to the lattice among tubulins at the plus end of MTs, whereas its C-terminal domain can associate with a wide array of proteins. The two domains operate in concert to accelerate MT dynamics (Rogers et al., 2002; Vitre et al., 2008; Maurer et al., 2014) and recruit various +TIP effector proteins that function at the plus end. In live cell imaging, fluorescent EB1 exhibits a comet pattern seemingly leading the plus end of nascent growing MTs, where tubulins transition from the GTP state to the GDP state (Maurer et al., 2012; Zanic et al., 2009; reviewed by Gardner et al., 2013). As such fluorescent EB1 has been commonly used to report newly generated growing MTs (Piehl et al., 2004; Matov et al., 2010). However, binding to the GDP zone increases in a number of scenarios (e.g. Tirnauer et al., 2002; Goldspink et al., 2013; Tortosa et al., 213; Sayas and Avila, 2014). What causes the switch remains uncertain.

Using EB1-NG as a reporter, we captured in real time unexpected changes in EB1-NG patterns and MT dynamics signaled through H^+^ and Na^+^. The remarkable sensitivity and the distinct responses in wild type (WT) cells and mutants shed critical insight on the divergence of the MT system, pH regulated processes and the vulnerability of organisms subjected to environmental stresses.

## Results

### EB1-NG reports remarkable sensitivity of the MT system in Chlamydomonas

The MTOC in *Chlamydomonas* cells at interphase is comprised of two pairs of basal bodies (BBs) (Figure 1a, top panel). The mother BB nucleates the assembly of the axoneme, a MT-based scaffold that drives the rhythmic beating of the flagellum. Four rootlet MT bundles (thick lines) arrange in a cruciate pattern positioning BBs at the apical end and the other organelles (Mittelmeier et al., 2011; Picariello et al., 2014). These MT bundles consisting of more than two acetylated stable MTs. In contrast, cortical MTs (thin lines) are singular (Horst et al., 1999) and highly dynamic (Harris et al., 2016). Under widefield fluorescence microscopy EB1-NG expressed at the level of endogenous EB1 from a genomic construct does not reveal stable MTs except the flagellar tip where plus ends of axonemal MTs undergo turnover continuously (Harris et al., 2016). In addition, plus ends of growing cortical MTs appears like the typical comets observed in other eukaryotic cells. Comets emerged from four spots underneath flagella, corresponding to BBs (bottom panel) (Pedersen et al., 2003). As shown from the top, side and rear views of cells, comets of nascent cortical MTs travel along the contour of the cell body toward the posterior pole (Figure 1b, white arrowhead; Supplemental video 1-1). Near the pole comets vanish presumably when MTs stop growing or switch to the shrinking phase. The pattern appears similar in cells resuspended in the commonly used Tris-Acetate-Phosphate (TAP) culture medium or 10 mM HEPES buffer (with 5 mM Na^+^).

**Figure 1.**
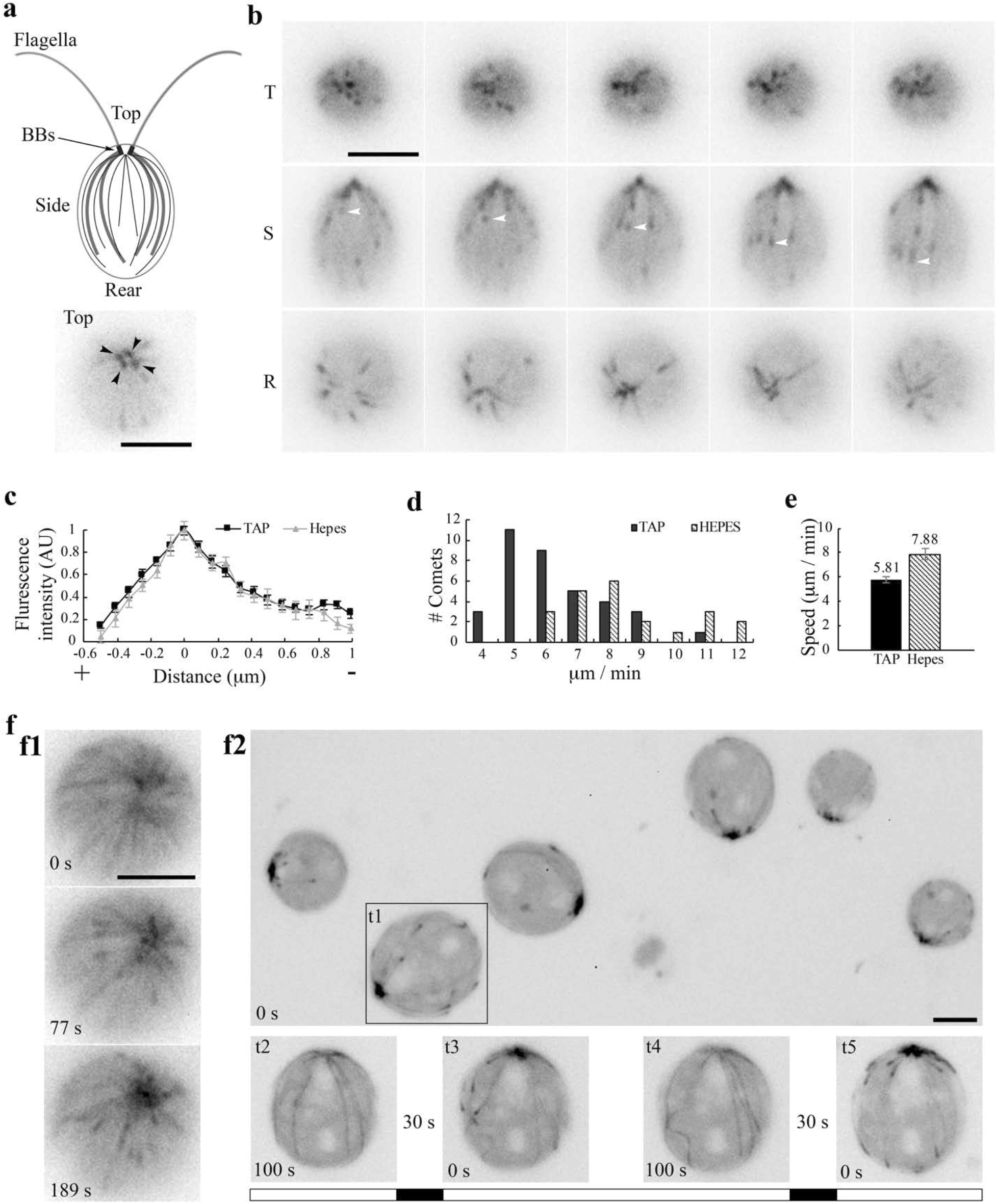
EB1 in *Chlamydomonas*. (**a**) A schematic picture depicting flagella and the MT network in the cell body (top panel). Black dots, basal bodies (BB). Thick lines, four stable rootlet microtubule bundles. Thin lines in the cell body, the dynamic cortical MTs. A top view of EB1-NG transgenic cells reveal a pattern that resembles 4 BBs (bottom panel). (**b**) Time-lap fluorescent images were taken 10 sec apart from the top (T), side (S) and rear (R) of cells resuspended in the TAP culture medium. EB1-NG appeared like typical comets (arrowheads), emerging from the BB area, coursing along the contour of the cell body and then vanishing as approaching the rear end. The frame rate is 1 frame/sec. (**c**) Normalized line scans along the length of MT plus ends showed a similar EB1 intensity profile in the TAP medium (n = 18 comets from 6 cells) and the Na^+^/HEPES buffer (n=11 comets from 3 cells). The position with peak intensity was designated as 0. The value was negative toward plus end; positive toward BBs. AU, arbitrary unit of fluorescence intensity. (**d**) The distribution and (**e**) the mean and the SEM of EB1 comet speed in the TAP medium (n = 36 comets from 6 cells in 6 recordings) and 5 mM Na^+^/HEPES buffer (n = 22 comets from 3 cells in 3 recording) are significant different (Mann-Whitney U test, P < 0.001). (**f**) Altered MT patterns during fluorescence microscopy. The EB1 comet pattern occasionally switched to a bird cage pattern (**f1**). Comets returned while the bird cage receded in ~ 1 min. In flattened cells that were compressed by the cover slip gradually, both MTs and comets became explicit (**f2,** top panel). Comets disappeared after ~ 100 sec (bottom panel, t2), but returned after illumination was switched off for 30 sec (t3). The process was repeatable after another 100 sec illumination and then another light off period (t4 and t5). The alternate white and black bars illustrate the scheme of alternate illumination and dark periods. Scale bars, 5 µm.

The birth of new comets from BBs appeared stochastic. We did not measure the birth rates, hindered by substantial fluctuations and the narrow apical area. Instead we analyzed comet length and speed from the side view. Line scans along the lengths of comets show the typical feature of EB1 comets - the brightest spot corresponds to the area where tubulins are primarily at the transitional state, slightly behind the leading edge of plus ends with GTP-tubulins (Figure 1c). The distribution of comet speeds shows that MT growth rates varies nearly two folds (Figure 1d). The dataset from cells in the TAP medium (black bars) skews toward the slow end relative to the Na^+^/HEPES dataset (hatched bars). The average velocities are signifantly different (Mann-Whitney U test, p < 0.001), at 5.8 ± 0.26 and 7.9 ± 0.42 µm/sec respectively (Figure 1e), which are within the normal range measured in diverse eukaryotic cells (Harris et al., 2016).

Curiously, in some long recordings, comets suddenly gave way to a bird cage-like pattern (Figure 1f1, top panel; Supplemental video 1-2) as if all cortical MTs were revealed by anti-tubulin immunofluorescence (Horst et al., 1999; Dymek et al., 2006). Comets returned automatically after ~1 min (middle and bottom panels). This unpredictability suggests that this pattern is caused by fluctuated intracellular conditions. When cells gradually flattened by the coverslip, MTs also became visible as a broken bird cage with comets (1f2, top panel, t1; Supplemental video 1-3). However, comets disappeared after ~ 100 seconds (bottom panel, t2). Interestingly, after a 30-second dark period, comets returned upon excitation light was switched back (t3). Simply alternating the dark and light period replicated the disappearance and return of comets (t4 and t5). These observations demonstrate that *Chlamydomonas* MT system is highly mercurial; and suggests that excitation illumination creates a condition that is unfavorable for MT dynamics, but is reversed in the dark. As illumination opens channelrhodopsins that conduct a number of cations and Cl^-^ (Nigel et al., 2002; 2003; reviewed by Hegemann and Berthold, 2009), we hypothesize that fluctuations of electrolyte concentrations modulate the MT system in *Chlamydomonas*. Considering the light sensitivity, we elected to use wide field microscopy and minimal light intensity to test this.

### Sequential changes in the MT system elicited by a short HA pulse and subsequent wash

We first used the well-defined pH shock, recording EB1-NG signals in cells exposed to HA in two complementary devices, perfusion chamber and diffusion chamber (Figure 2a). Recording of events in perfusion chambers started immediately before injection. Similar to pH shock, all cells were subjected to a swift change of environments as the injected solution was immediately pulled across the chamber by filter papers placed at the opposite end (Wheeler et al., 2008). However, the initial period was not decipherable due to flowing of injected fluid and unattached cells. For the diffusion chamber encircled by Vaseline, cells were placed at one side, underneath the coverslip and objective lens (right panel). Recording started after 100 mM HA was injected into the opposite side and cells stopped flowing. This design that decelerated the acidification process, captured the events as HA diffused across the field and into cells that were being recorded. However, the precise exposure time was uncertain; HA concentrations increased gradually, deflagellation was less effective; and not all cells were acidified to the same degree at the same time. Also the enclosed chamber cannot be washed.

Following the injection of 20 mM HA/TAP into the perfusion chamber, all comets vanished in the first discernable image taken at ~ 90 sec (Figure 2, b1-b2). After wash with 10 mM pH7 HEPES, comets re-appeared at the BB area within ~ 45 sec and MT dynamics resumed (b3-b5). Thus HA exposure elicits the disappearance of EB patterns either by perturbing EB-MT interplays or causing cortical MTs to pause or disassemble. New dynamic MTs re-form rapidly after HA is washed away.

Diffusion chambers in which the acidification process occurs gradually allow us to capture another unexpected phenomenon before comets vanish. A broken bird cage pattern with a few MTs (Figure 2c, left and middle panel, black arrowhead) and comets (white arrowheads) appeared before the disappearance at a time designated as 78 second (right panel). To decipher the disappearance, we analyzed digitally enhanced recordings (Figure 2d, top panels). As shown in two representative cells, overall EB1 signals were fading with time, which could be due to photobleaching, pH sensitivity of fluorescent proteins, disassociation of EB1 or system-wide MT disassembly. Some MTs clearly underwent endwise resorption (arrowheads). Kymographs tracking plus ends of prominent MTs in two rare still cells show that, as expected, comets disappeared before the resorption of the respective MT. Measurements of the slopes show that the initial shortening speeds for these two events are 8 and 13 µm/min but decelerate toward the BB area. These speeds revealed by EB1-NG are in line with the averaged 10±3 µm/min shortening speed of cortical MTs in cultured tobacco cells (Dixit and Cyr, 2004), faster than kinesin-13 catalyzed shortening (Helenius et al., 2006), and slower than the maximal shortening speed (30 µm/min) of MTs assembled from purified tubulins (O’Brien et al., 1997). Statistics of shortening speed was not analyzed because of few motionless cells and few shortening MTs with a definitive plus end. Tubulin reporters will be more appropriate for shortening analysis.

**Figure 2.**
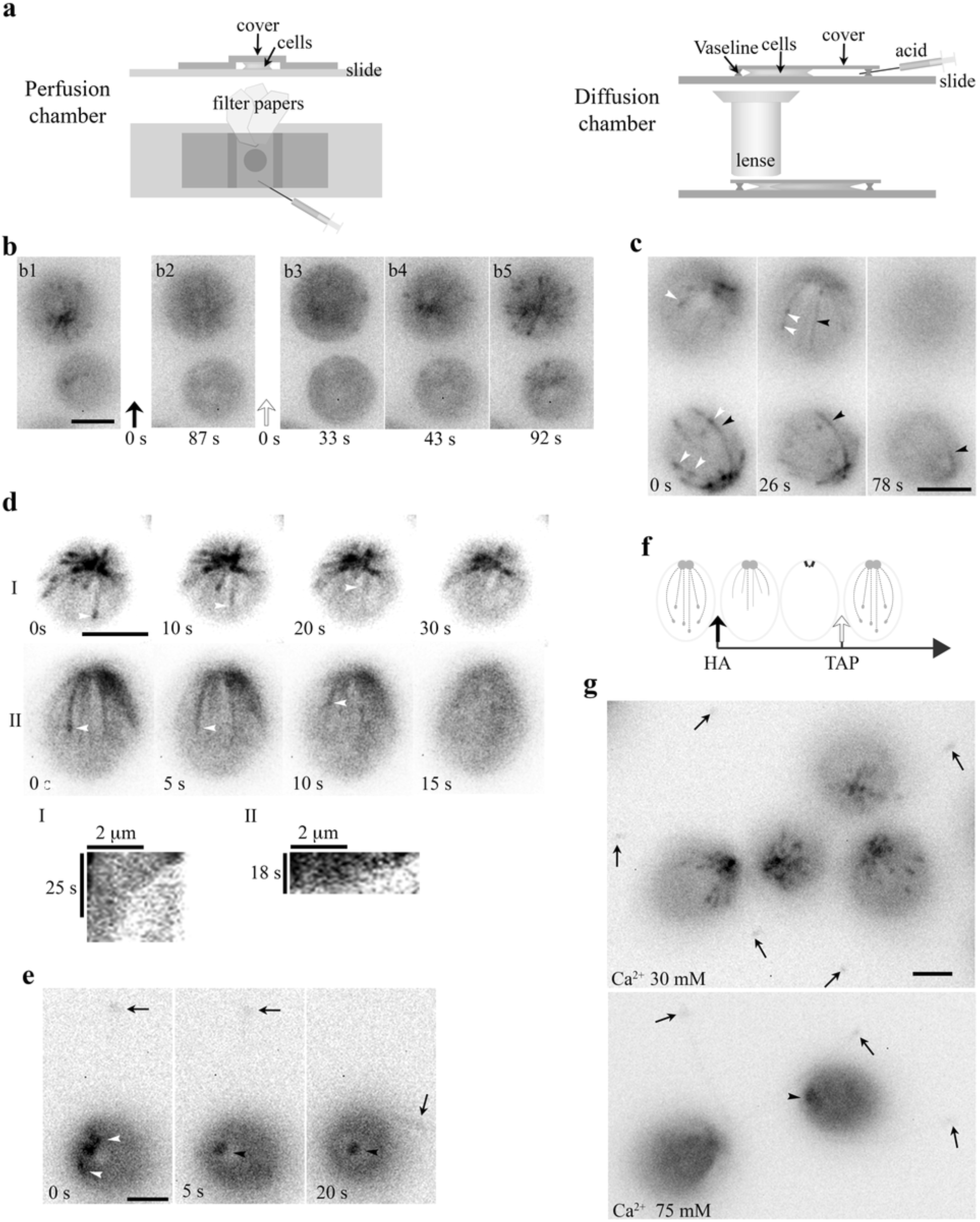
An HA pulse elicited swift sequential changes in the MT system. **(a)** Schematics depicting an open-ended perfusion chamber (left panels) and a diffusion chamber (right panels) for capturing the HA-induced rapid changes. **(b)** A 10- µl aliquot of cells resuspended in the TAP medium was placed in a perfusion chamber. The images (**b1, 2**) were captured before and after perfusion with 20 mM HA/TAP (pH 4.5, t=0, black arrow). The following recordings (**b3-5**) captured the events right after the TAP medium (pH 7) was injected to wash away HA (t=0, clear arrow). B3 is the first clear image after fluid and cells stopped flowing. Comets already disappeared within 87 seconds after HA perfusion. They started emerging 43 sec after wash. **(c)** The process preceding HA-induced disappearance of EB1 comets in diffusion chambers. A 40- µl aliquot of cells resuspended in HEPES was placed in a diffusion chamber encircled by Vaseline, under the coverslip and an objective lens. HA was injected to the other side of the chamber and diffused toward cells that were being imaged. During the gradual acidification process, both comets (white arrowheads) and shank binding MTs (black arrowheads) were evident first and then both patterns vanished. (**d**) Time lapse images and kymographs revealed endwise resorption of EB1-decorated MTs (white arrowheads). (**e**) Comets (white arrowheads) in the cell body vanished first before the excision of flagella (arrows). Following deflagellation, EB1 diffused away from the tip. EB1 signals remained at BBs but was static (black arrowhead). (**f**) A schematic depicting sequential changes in MTs upon exposure to HA and a subsequent wash with the TAP medium. Dotted lines with a comet, growing MTs. Dotted lines alone, shrinking MTs. Solid lines, shrinking MTs with EB1 shank binding. (**g**) Effects of [Ca^2+^]_ex_ on MTs. In cells resuspended in 30 mM [Ca^2+^]_ex_, flagella remained attached (arrows in left panel), while comets were vibrant. In the 75 mM [Ca^2+^]_ex_ group, comets disappeared and flagella were amputated (arrows in right panel). Static EB1 signals remained at BBs (arrowhead). Scale bars, 5 µm.

As for cells with EB1 signals detected in the cell body and flagella (Figure 2e) simultaneously, comets (white arrowheads in left panel) vanished first (middle panel) before flagella (arrow) were amputated (right panel). EB1 signals remained at the BB area but were static (black arrowhead). Contrary to deflagellation within seconds upon HA perfusion (Wheeler et al., 2008), the deflagellation in the diffusion chamber takes more than one minute due to gradual acidification. Thus when cells are exposed to HA, shank binding increases, comets disappear, endwise resorption becomes evident and then flagella become amputated. The sequential events occurring in the diffusion chamber are summarized in Figure 2f.

Lowering intracellular pH elicits Ca^2+^ influx, whereas Ca^2+^ prevents MT formation and promotes MT disassembly (Weisenberg, 1972; O’Brien et al., 1997). To differentiate whether HA-induced changes are due to H^+^ or Ca^2+^, we raised [Ca^2+^]_in_ without adding HA. Calcium ionophore A23187 did not trigger deflagellation or evident changes in the MT system in our hands. This is not surprising since A23187 cannot elicit consistent effects in *Chlamydomonas* (Bloodgood and Levin, 1983). So we simply raised [Ca^2+^]_ex_. Perfusions of either HA or CaCl_2_ solution elicit Ca^2+^ influx, leading to deflagellation, although the latter is less efficient (Wheeler et al., 2008). WT cells resuspended in 30 mM Ca^2+^/HEPES appeared agitated, suggesting entry of Ca^2+^ (Figure 2g, top panel). However, flagella remained attached (arrows) and comet activity was robust. When cells were resuspended in 75 mM Ca^2+^/HEPES solution (right panel), cells shed flagella (arrows). Comets already disappeared, while static EB1 signals remained at the BB area (arrowhead). Thus the outcomes elicited by high [Ca^2+^]_ex_ and HA are similar. The experiments testing the effects of Ca^2+^ and hypertonicity will be described in later sessions.

### Formation of cold-resistant thick MT fibers in the recovery phase after HA bath

*Chlamydomonas* expresses Na^+^/H^+^ exchangers (Pittman et al., 2009), as well as various channels and pumps at different locations (Fujiu et al., 2011; Taylor et al., 2012) to maintain electrolyte homeostasis. We reasoned that altering HA treatment might also change other cations such as Na^+^ or K^+^. To test this, we extended HA exposure - resuspending cells in pH3, 10 mM HA/double distilled water (ddw) for 5 mins. As expected, EB1 patterns were absent except for the static signal at the BB area (Figure 3a, left panel), mimicking the final outcome of HA exposure in the diffusion chamber (Figure 2e). In contrast, resuspension in pH3, 10 mM HCl did not alter comet activity. Thus, as with deflagellation, HA-elicited changes in cortical MTs are due to intracellular rather than extracellular acidification. Images were taken as cells were washed with Na^+^/HEPES in a perfusion chamber. After washing, dynamic EB1 signals emerged at the BB area after ~50 sec (Figure 3a, cell I in right panel). Interestingly, nascent MTs were not adorned with a typical comet. Compared to the bird cage pattern, they appeared thicker, fewer and nearly uniformly decorated, as if plus end tracking EB1 stayed behind growing MTs. In cells recorded at a later period (cell II and III, between 60 – 159 sec; Supplemental video 3), MT growth decelerated, especially between 109-159 sec. In cells recorded after 180 sec, all growth stopped (Figure 3b, left panel). This is further illustrated by two nearly identical images taken 20 sec apart of two representative cells (cell I and II, right panels). Notably, some static fibers split, or had more than one comet aligned in tandem (arrowheads). Taken together, these observations strongly suggest that nascent MTs generated after HA bath and wash are abnormal, perhaps with a propensity to nucleate ectopically, branching or growing on top of the other piggy-backing as bundles. Kymograms comparing three representative MTs (Figure 3c, top panels) with growing MTs in the untreated control cells (bottom panels) confirm that EB1 signals extend further toward the BB area in cells recovering from HA bath. MT growth rates, shown by the slopes, fluctuate but were mostly slower than those in untreated control cells. Overall the rates decline until MT growth stops. Thus contrary to thin, dim, transient MT fibers in the acidification phase (Figure 2), MTs formed in the recovery phase after HA bath were thick, short, long-lived and bright.

MTs are cold labile. Cold treatment induces MT endwise resorption *in vitro* (Muller-Reichert et al., 1998). To test the stability of these thick MTs formed in the recovery phase, glass slides with a droplet of cells after HA bath and wash were placed on ice for 3 min. Images were taken using the microscope at the room temperature (RT) immediately, about 20 seconds after slides were removed from ice. Unexpectedly, most cells were imaged from the apical end and the focal planes drifted continuously, indicating that cells oriented toward the objective lens and floated gradually during this warm up period. As shown in two representative cells, EB1-decorated MTs after HA bath and wash remained after the removal from ice amidst the drift of focal planes (Figure 3d, top panels). Thus the thick static MTs formed after HA bath and wash are cold stable. The pattern remained for the subsequent 70-second recording period. In contrast, for the control without HA bath, EB1 patterns were undetectable initially (bottom panels), indicating cold lability. Dynamic EB1 signals gradually re-emerged at BBs after 30 second (white arrowheads).

**Figure 3.**
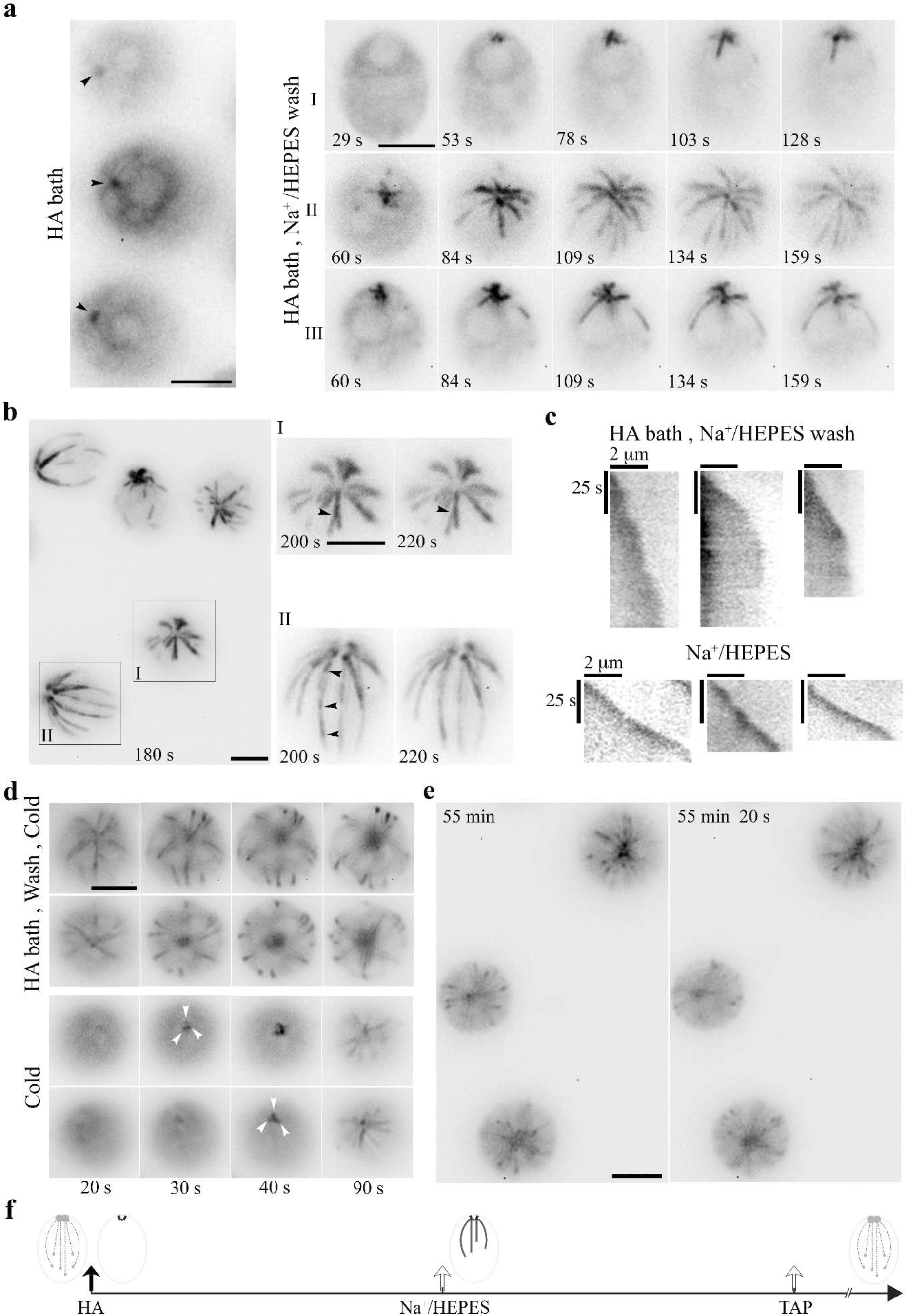
HA bath and a subsequent wash induced long-lived yet reversible changes to the MT system. (**a**) Static EB1 signals remained at BBs after cells were resuspended in 10 mM pH3 HA for 5 min (left panel). After replacing HA with the HEPES buffer, EB1 signal at the BB area intensified within 1 min (black arrowheads). But newly formed MTs were thick and prominent, lacking the typical comet (right panel, cell I). In cells recorded 60 sec after wash (cell II and III), MT elongation slowed down gradually. (**b**) In cells imaged ~180 sec after wash, EB1-decorated MTs in all cells stopped growing (top panel), as highlighted in two additional images of two representative cells captured 20 sec apart. In addition, a MT fiber in cell I split into two (arrowhead), while a fiber in cell II had multiple comets aligned in tandem (arrowheads), as if new MTs nucleated or grew on older ones. (**c**) Kymograms comparing the growth of three representative MTs in cells pretreated with 5-min HA bath and then the wash buffer (top panels) and in control cells in the HEPES wash buffer (lower panels). Comets in the latter manifested as intense spot at the plus end. The sharper slopes in the former indicated slower growth and eventually unchanged as the growth paused. (**d**) HA - induced long-lived MTs formed after HA bath and wash were cold resistant (top panels). As shown in two representative cells, frozen MTs remained in the image captured 20 sec after 3 min on ice. The changes in subsequent images were due to drift of focal planes as cells were floating gradually as warming up. In control cells without previous HA exposure (bottom panels), EB1 signals were absent initially (20 sec), consistent with cold lability. They gradually emerged at the BB area (white arrowheads) afterwards. (**e**) Although the MT system froze within minutes after HA bath and HEPES wash (t=0 min), comets resumed 55 mins in cells recovered in TAP media as shown in two images captured 20 sec apart. Scale bars, 5 µm. (**f**) A schematic summarizing the sequence of MT changes induced by HA bath and wash.

To learn if frozen thick MTs after HA bath and wash were reversible, we continued imaging cells in the recovery phase. To prepare for the long recovery, wash buffer was replaced with TAP media. Thawing signs emerged gradually. Compared to frozen thick MTs (Figure 3e, left panel, black arrowheads), at 55 mins post wash, comet (white arrowheads in right panel) activity was vibrant as if the freeze had never occurred. The sequential events occuring after HA bath and wash are summarized in Figure 3f.

### Testing HA-induced phenomena by using a tubulin mutant and by reducing HA concentrations

In an attempt to decelerate pH-induced resorption in the acidification phase, we took advantage of a tubulin mutant, *tub2*. A missense mutation near the colchicine binding site in β-tubulin increases MT stability since *tub2* cells are colchicine-resistant, and have more acetylated MTs (Schibler and Huang, 1991). In the EB1-NG transgenic *tub2* cells, the comet pattern (Figure 4a, top panels) appeared indistinguishable from that in WT transgenic cells (Figure 1b). Thus our assay is not sensitive enough to report increased MT stability. Interestingly though, in the image taken immediately after resuspension in 10 mM HA, the bird cage pattern with a few comet-like spots already formed in every *tub2* cell (bottom panels). The pattern was rather stable, although some MTs seemed to be out of focus intermittently (bottom right panel), suggesting detachment from the plasma membrane. As shown in two representative cells, similar frozen bird cage pattern remained even after 5-min HA bath (Figure 4b; Supplemental Figure 4-1).

**Figure 4.**
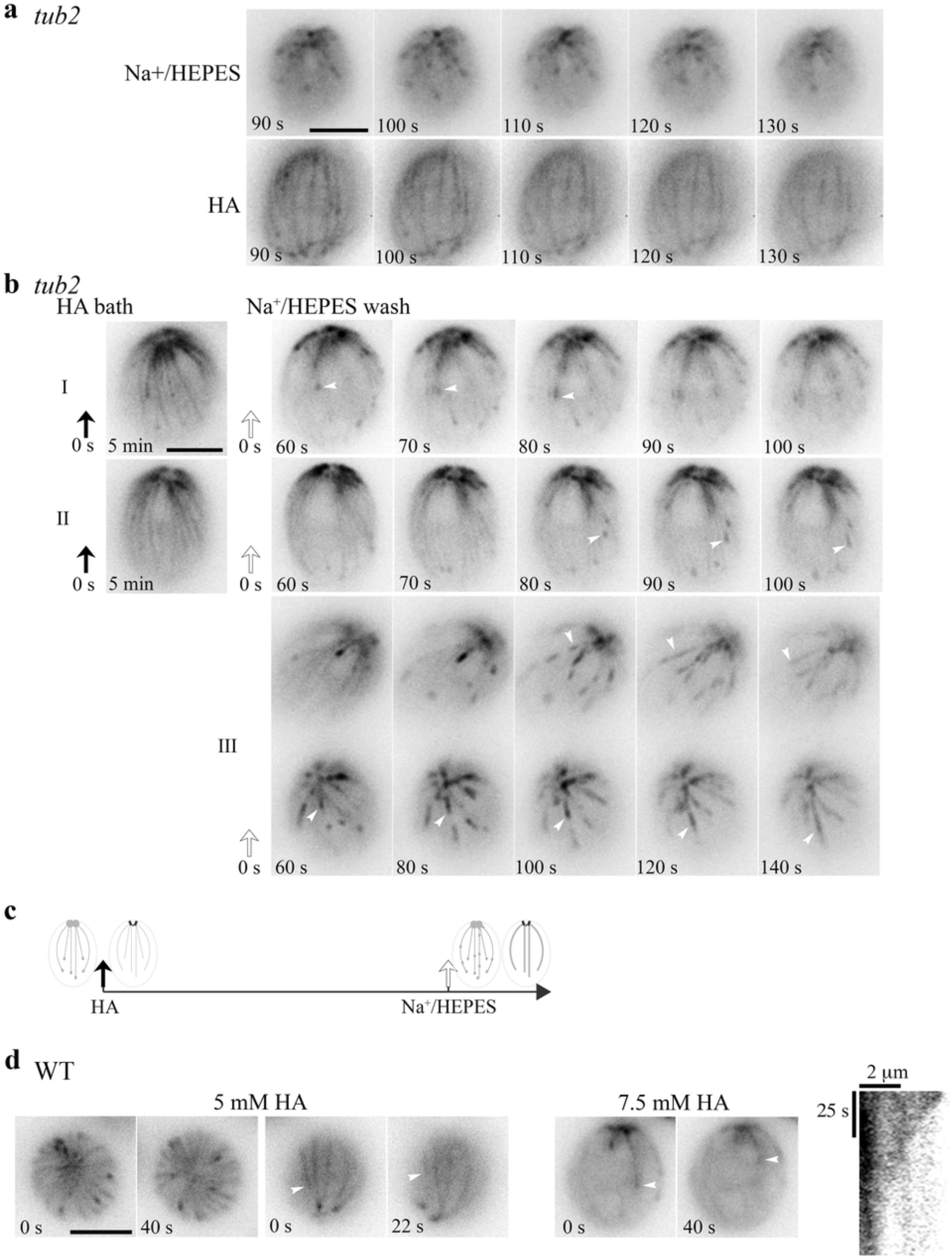
Tempered HA-induced changes by a β-tubulin mutation (**a-c**) and by reduced HA concentrations (**d**). (**a**) Time-lapse images at 10 sec intervals showed normal comets in *tub2* cells resuspended in 5 mM pH 7.4 Na^+^/HEPES (top panels). In contrast, a bird cage pattern with fine MT fibers was present in *tub2* cells resuspended in 10 mM, pH 3 HA (bottom panels). Most MTs froze but some appeared in and out of focus. (**b**) Frozen bird cage remains after 5-min HA bath as shown in two representative cells (left panel). After wash, comets (white arrowheads) reemerged at the plus end of existing fine MTs, instead of the BB area, while the bird cage pattern was fading. MTs were growing but comets were lengthening and the pace was very slow. In videos recorded only after wash, the recovery was faster. Some comets already emerged from the BB area. Some moved along old MTs and lengthened (panel III, white arrowheads). (**c**) A schematic summary of HA-induced changes in *tub2*. (**d**) Concentration-dependent effects of HA on MTs in WT cells. WT cells resuspended in 5 mM HA exhibited a dynamic bird cage pattern with dynamic comets (left, top and side view). Shrinking MTs were visible in the two side view images taken 22 sec apart (white arrowhead). Most patterns were absent in cells resuspended in 7.5 mM HA, except a few shrinking MTs (right). A kymograph revealed the endwise resorption. Scale bars, 5 µm.

After wash, comets returned (panel I and II) and lengthened as in WT cells. However, comets (white arrowheads) appeared at the plus end of existing MTs first rather than at BBs where nascent MTs emerged. The bird cage pattern gradually faded concomitantly. For images taken 60 seconds and later after wash, comets formed and moved along existing MTs (Panel III, Supplemental video 4-1). This strengthens the interpreted bundling propensity of MTs in WT cells recovered from HA bath (Figure 3b). These collective results indicate that the mutation in *tub2* hinders endwise resorption elicited by HA exposure, as such the bird cage pattern with cortical MTs uniformly decorated with EB1 is not vanishing; the other responses in *tub2* and WT cells are similar; and old MTs resorb to make room for new ones to maintain the number of cortical MTs. The HA responses of *tub2* cells are summarized in Figure 4c.

We further tested whether we could replicate the bird cage pattern in WT cells with less concentrated HA. As shown in two top view images taken 40 seconds apart immediately after resuspension in 5 mM HA (Figure 4d, left panel; Supplemental video 4-2), all WT cells had motile flagella and had a dynamic, rather than still, bird cage pattern in which MTs shrink or grow with a comet, as if the entire MT system were revealed by fluorescent tubulins and fluorescent EB1 simultaneously. The side view images recorded 22 seconds apart revealed endwise resorption of a shrinking MT (arrowhead). Only a few resorbing MTs were captured in cells resuspended in 7.5 mM HA (Figure 4d, right panel). This still cell allows us to plot the kymograph, which shows a tapered endwise resorption with the initial shortening speed of 4 µm/min. All MT patterns and comets disappeared in cells resuspended in 10 mM HA. Therefore, low [HA] increases EB1 shank-binding, rendering the bird-cage pattern. As HA concentrations increase, MTs stop growing and comets are lost. As resorption continues, perhaps even at a hastening pace, comet and bird cage patterns vanish. The 5 mM HA experiment consistently replicates the light-induced sporadic transient appearance of the bird cage pattern in WT cells (Figure 1, f1). Changes elicited by 7.5 mM HA partially mimics HA-induced responses in diffusion chambers (Figure 2b-e).

### The long-lived EB1-decorated MTs are due to the rise of intracellular [Na^+^] but not [K^+^]

To identify the ion causing the formation of static thick EB1-decorated MT bundles in cells recovering from HA bath, HA-bathed cells were washed with different solutions. Interestingly, as shown by time-lapse images separated by a 10-second interval, EB1-decorated MT fibers formed only if the wash solution contained Na^+^, such as the NaOH-buffered HEPES (5 mM Na^+^) or 5 mM NaCl in ddw (Figure 5a, top two panels). On the other hand, comets resumed profusely if the wash solution lacked Na^+^, such as the KOH-buffered HEPES buffer, 5 mM KCl in ddw or plain ddw (bottom three panels). Therefore, Na^+^ accounts for the reformation of thick, long-lived static MTs in the recovery phase of HA-bathed cells.

Since Na^+^ has low permeability compared to K^+^ (Ronkin and Buretz, 1960), we predict that Na^+^ ions from the wash buffer are entering the cytosol when Na^+^/H^+^ exchangers are removing H^+^. To test this, we resuspended WT cells in 10 mM, pH 8 EGTA buffered by NaOH (final ~21 mM Na^+^). Concentrated EGTA and EDTA are classical tools for creating permeable cell models perhaps by Ca^2+^ chelation or other mechanisms (Miller, 1979; Tazawa and Shimmen, 1983; Arikawa and Suzaki, 2002; Prachayasittiku et al., 2007). However, EGTA at this concentration does not influence polymerization of MTs from purified tubulins *in vitro* (Olmsted and Borisy, 1975). As expected, thick EB1-decorated MTs formed after cells were resuspended in Na^+^/EGTA for 5 min (Figure 5b, left panels). Thick MTs appeared static after 10 mins. In contrast, cells resuspended in KOH-buffered EGTA had vibrant comet activities (right panels). The static EB1-binding MTs in Na^+^/EGTA treated cells were also cold-resistant.

As K^+^/ EGTA did not affect comet activity, we depleted [Ca^2+^]_ex_ with K^+^/ EGTA rather than Na^+^/ EGTA to test if Ca^2+^ is required for the changes elicited by10 mM HA. After 5 min in K^+^/ EGTA, 10 mM HA resuspension still elicited the bird cage pattern in *tub2* cells, and caused comets to vanish in WT cells. However, cells quickly burst, indicating perturbed plasma membrane. Thus HA-elicited changes in MTs do not require extracellular Ca^2+^.

Despite low Na^+^ permeability of the plasma member, we further tested if boosting Na^+^ in HEPES buffer to 55 mM with NaCl was sufficient to change EB1 patterns in WT transgenic cells. Indeed, compared to the growing MTs with typical comets in the control cells in the 5 mM group, in the 55 mM group comets were evidently but MTs were growing (panel I). This is reflected by the modest tapering of comet intensity in the linescan plot (panel II). Therefore, high [Na^+^]_ex_ still could raise [Na^+^]_in_, increasing the time EB1 spent at the plus end and thus comet lengths. Despite the long comets, the distribution of comet velocity (panel III) and the average velocity (panel IV) show that [Na^+^]_in_ at this level only slightly reduces MT growth rate (two-tailed p-value = 0.0497, < 0.05). Thus both comet length and MT growth are sensitive to [Na^+^], albeit the former has a higher sensitivity.

To test the effect of hypertonicity, WT transgenic cells were resuspended in 150 and 200 mM [Na^+^]_ex_ for 5 min before imaging. At these concentrations, flagellar motility ceased. As in 75 mM [Ca^2+^] solution (Figure 2g), most comets vanished, although static EB1 signals remained at the BB area (Figure 5d). However, some cells still contain thick fibers (top panel), a signature of Na^+^-induced changes. Together, these results show that low Na^+^ and high Na^+^ elicit disparate MT responses. The modest increase of [Na^+^]_in_ from 55 mM [Na^+^]_ex_ alone promotes long comets but is insufficient to stop MT growth. The fact that both responses are triggered by HA bath followed by 5 mM Na^+^/HEPES wash or by 21 mM Na+/EGTA strongly suggests [Na^+^]_in_ rises further to a level under 21 mM due to H^+^/Na^+^ exchange or passive diffusion through EGTA-permeablized membrane. However, most EB1 signal vanished, suggesting MT disassembly at the concentrations that deem hypertonic for *Chlamydomonas* either via [Na^+^]_ex_ and/or [Ca^2+^]_ex_.

**Figure 5.**
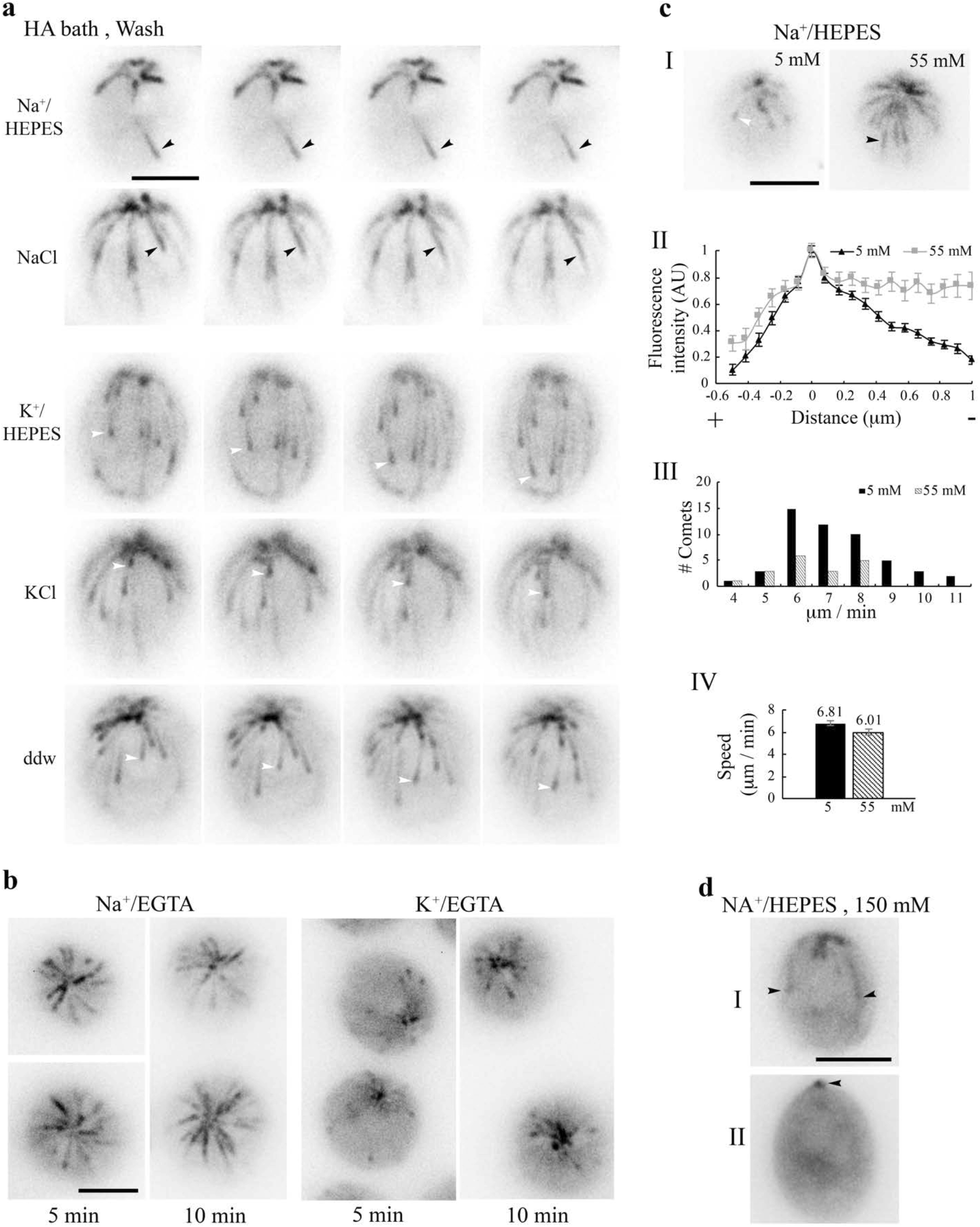
Na^+^-dependent changes of the MT system. (**a**) MTs in cells were largely frozen after 5-min 10 mM pH3 HA bath and 3 min in the wash solution, such as 5 mM pH7.4 Na^+^/HEPES buffer or 5 mM NaCl solution (black arrowheads). In contrast, growing MTs with a comet (white arrowheads) returned if the wash buffer lacked Na^+^, such as 5 mM K^+^/HEPES buffer, 5 mM KCl solution, or the double distilled water (ddw). (**b**) Thick MTs in cells resuspended in 21 mN Na^+^/EGTA for 5 min or 10 min (left panel), contrary to comets in cells in 21 mM K^+^/ EGTA (right panel). Thick MTs were still growing after 5 min incubation but static after 10 min incubation (**c**) High [Na^+^]_ex_, without preexposure to HA, was sufficient to alter comet patterns. Contrary to typical comets in cells resuspended in the HEPES buffer with 5 mM Na^+^, long comets were thick in cells resuspended in 55 mM Na^+^ for 5 min (panel I). Normalized linescans confirmed little tapered intensity (panel II, n=36 comets from 11 cells in 5 mM Na^+^; n=13 comets from 4 cells in 55 mM Na^+^). As shown in the range of speed (panel III) the long comets were moving, and the mean speeds of short and long comets were significantly different (panel IV, n=51 from 11 cells in 5 mM Na^+^; n=18 from 4 cells in 55 mM Na^+^) (P < 0.05). (**d**) Two representative cells after 5 min in 150 mM Na^+^/HEPES. Some cells still retained a few thick MTs (cell I). Some only had static EB1 signals at the BB area (cell II, arrowhead). Scale bars, 5 µm.

## Discussion

Expression of EB1-NG as a reporter loosens the bottle neck posed by autofluorescence of *Chlamydomonas* and unleashes the potential of this MT model system. Contrary to the perceived stability of the MT system in typical interphase animal cells (Lieuvin et al., 1994), EB1-NG reports in real time the mercurial changes of algal MT system that are elicited by excitation illumination, compression, H^+^ and Na^+^. We summarize the changes and discuss possible underlying mechanisms; and the new insight on the MT system, plant salinity responses and additional concerns of environmental stresses.

### Changes elicited by H^+^

The changes of algal MT system elicited by intracellular acidification are swift, stunning and novel (Figure 2b-e, 3a and 4). Among heightened shank binding, diminished comet activity, paused MT growths, and MT shortening, the bird cage pattern of shank binding is the most sensitive, elicited reliably by 5 mM HA (Figure 4d). They are unlikely signaled through cell death pathways, since these changes are reversible, even after 5 min HA bath (Figure 2b, 3). The recovery occurs within a minute, either completely (Figure 2b) or protractedly (Figure 3e), depending on exposure and wash procedures. Extracellular Ca^2+^ is not required for the HA-induced changes, since they still occur after K^+^/EGTA treatment. However, we cannot rule out the involvement of Ca^2+^ released from intracellular storages and other signaling pathways.

Although the pH of [HA]_ex_ that triggers these changes is ~3, we expect that the resulting intracellular pH is close to or higher than 6.3. HA-induced changes in diffusion chambers appear before deflagellation (Figure 2e) that occurs at pH 6.3 (Wheeler et al., 2008; Braun and Hegemann, 1999). Consistent with this, the bird cage pattern could be transiently triggered merely by illumination that may open H^+^-selective channelrhodopsin (Figure 1f1). Conversely, comets return to compressed cells when illumination is turned off for 30 seconds (Figure 1f2); or return within ~ 45 sec after HA is washed away (Figure 2b). These observations strongly suggest that a slight imbalance of pH homeostasis is sufficient to elicit changes in algal MT system.

Although pH affects proteins’ ionization and thus their functions and protein-protein interactions in general (Hepler, 2016), we speculate that the MT system is particularly sensitive to declining pH because of the acidic pI of tubulins and EB1. For example, the respective pI of *Chlamydomonas* α-tubulin, β-tubulin and EB1 is 5.01, 4.82, and 5.7. A decrease of pH from the resting level will make these proteins less negatively charged, especially at their C-terminal acidic tails that are central to MT-accessory protein interplays and the targets of various post-translational modifications (reviewed by Song and Brady, 2014; Buey et al., 2011; Rovini et al., 2013). The resulting decreased repulsion could explain increased affinity of EB1 to GDP-tubulin, leading to nearly immediate appearance of the bird cage pattern in *tub2* cells (Figure 4a-b) or in WT cells exposed to 5 mM HA (Figure 4d). As pH descends further, additional changes in protein conformation may inhibit the growth of MTs and EB1 binding to plus ends, leading to comet reduction or ultimate disappearance.

These HA-induced changes explain long standing questions regarding pH variations. For example, the pollen tube tip has overlapping regions. The MT zone, in particular, lags behind the F-actin, Ca^2+^ and acid zone (Gibbon and Kropf, 1994; reviewed by Hepler, 2016). Likewise, a basic shift directs MT-supported fertilization processes of sea urchins, whereas depressing pH inhibits the processes and triggers MT disassembly (Schatten et al., 1985). This and an increase of 0.3-0.5 pH unit in mitosis inspired the pH clock hypothesis for cell cycle control (Gagliardi and Shain, 2013). In line with this, EB1 preferentially binds to MT plus ends in arrested mitotic phase extract of *Xenopus* oocytes, but uniformly decorates MTs in interphase extract (Tirnauer et al., 2002) perhaps with a lower pH, analogous to HA-induced bird cage pattern (Figure 4c). Given the role of EB1 in recruiting effector molecules and the swiftness of pH-induced changes in MTs and EB1 patterns, tuning pH may indeed control cell cycle at least for certain organisms.

### Changes elicited by Na^+^

The responses elicited by [Na^+^]_in_ and [H^+^]_in_ at low concentrations are distinct. Contrary to instant appearance of H^+^-elicited bird cage pattern with fine, individual MTs (Figure 4d), as Na^+^ continues rising, comets lengthen; cortical MTs undergo ectopic nucleation, splitting, bundling, decelerate and stop eventually (Figure 3 and 5). Although it takes longer to elicit Na^+^ responses, this is likely due to low Na^+^ permeability. Likewise, Ca^2+^ permeability is tightly controlled. As such [Na^+^]_in_ and [Ca^2+^]_in_ cannot be adjusted as nimbly as [H^+^]_in_.

The degree of changes correlates with [Na^+^]_in_. These changes aggravating with time in 21 mM Na^+^/EGTA (Figure 5b) indicates that they are occurring before [Na^+^]_in_ reaches 21 mM, which is made possible by EGTA treatment. As EGTA chelates Ca^2+^, this further rules out the involvement of extracellular Ca^2+^. Similarly, Na^+^-dependent responses emerge within ~45 seconds once HA bath is replaced with various solutions containing only 5 mM Na^+^ (Figure 5a), suggesting accelerated rise of [Na^+^]_in_ due to the activity of Na^+^/ H^+^ exchangers (Pittman et al., 2009). On the other hand, by simply relying on limited passive diffusion through the normal plasma membrane, 55 mM [Na^+^]_ex_ is sufficient to change comet length but only slows down growth rate slightly (Figure 5c). Based on the incremented responses, rather than all-or-none responses to a threshold, we speculate that algal MT system is also sensitive to [Na^+^]_in_, perhaps in a linear manner. Contrary to [Na^+^]_ex_, raising [K^+^]_ex_ had no evident effect (Figure 5 a-c). This is reasonable, given high [K^+^]_in_, ~ 70 mM in *Chlamydomonas* cells (Malhotra, 1995). It highlights the selective sensitivity of the algal MT system to Na^+^ and rules out mere ionic effects. One interesting possibility is that Na^+^ binds to particular sites in algal tubulins, analogous to Ca^2+^ binding sites in mammalian tubulins (Solomon, 1977; Serrano et al., 1986).

### Common changes elicited by high extracellular HA, Na^+^ and Ca^2+^

EB1 signals largely vanish at 150 mM [Na^+^]_ex_ (Figure 5d), as in 10 mM HA (Figure 2c, 3a) and 75 mM Ca^2+^ (Figure 2g) except residual static signals at the BB area. They are likely caused by synergistic disassembly and paused new growth, and an immotile EB1 population underneath BBs respectively (Yan et al., 2006; Pedersen et al., 2003). The similar outcomes caused by distinct ions and obvious shrinkage of the cell body at even higher concentrations of Na^+^ and Ca^2+^ suggest that hypertonicity is involved. We envisage that high concentration responses could be caused by one cation exceeding a threshold concentration; and/or simultaneous rises of multiple electrolytes as H2O moves out of cells. Hypertonicity may evoke additional pathways.

The capture of endwise resorption only in low concentration conditions (Figure 2d and 4d) suggests that increased concentrations of these ions will heighten shortening-the incidence and/or speed. This is reminiscent to high Ca^2+^ effects. *In vitro*, Ca^2+^ blocks MT formation (Weisenberg, 1972), whereas 0.5-0.6 mM Ca^2+^ - in the absence of MAPs - could increase shortening incidence and accelerate shortening speed of MTs beyond 150 µm/min (Karr et al., 1980; O’Brien et al., 1997). Although pH shock and high [Ca^2+^]_ex_ only temporarily raise [Ca^2+^]_in_ up to 1 µM (Wheeler et al., 2008), the lower concentration may be sufficient to heighten shortening. However, this possibility is weaken by the same result despite EGTA treatment.

Other proteins and signaling pathways are involved in the electrolyte-elicited responses. Kinesin-13 that catalyzes MT disassembly at a lower speed (Helenius et al., 2006) becomes phosphorylated within 5 min following pH shock; and knockdown of kinesin-13 ameliorated the pH shock-induced reduction in MT lengths and numbers (Wang et al., 2013). Although EB1 patterns appear fully recovered 1 min after a pH pulse (Figure 2b), quantitative analysis and using similar reporters are needed to compare the results. Similarly, osmotic or salt stresses activate an atypical tubulin-kinase and phospholipase D to trigger disassembly or reorganization of plant MT system (Fujita et al., 2013; Dhonukshe et al., 2003). For yeast, sorbitol hypertonicity induces frozen MTs (Robertson and Hagan, 2008) and the recovery in 38 mins involves a stress-induced MAP kinase. Similar paradigms may be responsible for the resumption of MT dynamic in HA-bathed algae that takes ~55 mins (Figure 3f).

The electrolyte sensitivity of algal MT system is contrary to the perceived stable MT system in interphase mammalian cells (Lieuvin et al., 1994) that have 140 mM [Na^+^]_ex_, and 140 mM [K^+^]_in_ (reviewed by Pohl et al., 2013). Consistent with this, we cannot elicit any obvious changes of EB1-EGFP patterns in mammalian epithelial cells by compression or illumination (Matov et al., 2010). Likewise, both Na^+^ and K^+^ promote tubulin polymerization, with 160 mM Na^+^ as the optimal condition (Olmsted and Borisy, 1975). Different cation sensitivities could be due to sequence divergence. The other is the presence of different accessory proteins. MAPs obscure the cation sensitivity of mammalian MTs (Olmsted and Borisy, 1975; Wolff et al., 1996). Alternatively, signaling pathways or the capacity to maintain electrolyte homeostasis could differ. Thus while fundamental features of the MT system - likely dynamic instability and EB1 plus end tracking - are universal, electrolyte sensitivity and responses could diverge.

The sensitivity of algal MT system to Na^+^ is consistent with enlarged or clustered algal cells cultured in high salt media (Takouridis et al., 2015). This could be caused by anomalies in the MT-supported processes in the cell cycle, such as mitosis and trafficking-dependent release of hatching enzymes (Kubo et al., 2009). Yet this fresh water green alga has several strategies to adapt to salinity (Perrineau et al., 2014), such as glycerol production (Husic and Tolbert, 1986), switches in gene expression (Gao et al., 2016), sexual reproduction and mutations (Takouridis et al., 2015). Salinity adaptation and the incredible H^+^ and Na^+^ sensitivity of algal MT system that bears semblance to that in both animal and plant cells demand a fresh look at how environmental stresses affect diverse organisms.

One is ocean acidification by anthropogenic CO_2_ (Raven et al., 2005). Hindered shell formation from the resulting extracellular acidification with pH declining by merely 0.1 unit has rightfully raised much alarm (Hoegh-Guldberg, et al., 2007; Waldbusser et al., 2013; Fitzer et al., 2016). Similarly, 5 mM and 7.5 mM HA differ 0.09 pH unit based on the pK_a_ of HA and the Henderson-Hasselbalch equation, but the intracellular acidification from this slight difference triggers distinct changes of *Chlamydomonas* MT (Figure 4d). Notably, dictated by CO_2_ chemistry, the ratios of permeant CO_2_ and H_2_CO_3_ to non-permeant ionic forms will increase further as solutions acidify, aggravating intracellular acidification. Therefore, intracellular acidification could be equally, if not more, insidious to marine species that are not equipped to cope with this stress; and may poise to shape aqueous landscapes and drive evolution (Cannon et al., 1985).

The other is salt stress and osmostress caused by draught, which is exacerbating due to climate changes or improper agriculture practices. The H^+^-enhanced, Na^+^-dependent responses of algal MTs (Figure 3a, 5) - bundling, cold-resistance and growth inhibition - resemble salinity-elicited changes in plants (e.g. Dhonukshe et al., 2003; Shoji et al., 2006; Wang et al., 2007; Zhang et al., 2012; Fujita et al., 2013; reviewed by Hardam, 2013; Hashimoto, 2015; Oda, 2015). The similarity comports with their common cortical MTs and a great homology of their proteins in the MT system (Dymek and Smith, 2006; Pedersen et al., 2003; reviewe by Gardiner, 2013) except flagellar genes (Merchant et al., 2007). MT changes induced by various abiotic stresses, including salt, have prompted an interesting proposition that the MT system is an abiotic sensor of plant cells (Haswell and Verslues; 2015; Wang et al., 2011). Yet the involvement of signaling pathways and slow readouts in plant experiments question whether the changes of MTs are the consequence of salt stress instead. The EB1-reported, scaled responses that seem proportional to [Na^+^]_in_ and the speed of manifestations (Figure 4d and 5) in fresh water algae strengthen the possibility that plant MT system is an upstream player in the salinity signaling pathways, if not the very sensor. In line with this, channel-linked MTs are integral to osmolarity signaling transduction in mammalian osmosensory neurons (Prager-Khoutorsky et al., 2014). Using the experimental strategies developed in this study, it is possible to investigate quantitatively the diverged mechanisms of eukaryotic MT systems in sensing and responding to salt stresses.

## Materials and Methods

### *Chlamydomonas* Strains and Culture

The wild type strain CC-124 and a β-tubulin mutant *tub2-1* (formerly *col^R^4*) (Bolduc et al. 1988; Lee et al. 1990; Schibler et al. 1991) were from *Chlamydomonas* Resource Center (<u>http://www.chlamycollection.org/</u>). They were converted into EB1-NG transgenic strains as described (Harris et al. 2016). Cells were cultured in 300 ml pH7.0 standard tris/acetate/phosphate (TAP) liquid media with aeration at 25°C over a 14/10 light/dark cycle until reaching logarithmic phase of growth (5-10 X 10^6^ cells/ml) (Sivadas et al., 2012). All experiments were completed at least 2 hours before the onset of the dark period.

### Solutions

Glacial acetic acid (HA) was diluted with ddw to various concentrations ranging from 5 to 1000 mM. The 10 mM HCl solution was titrated to pH3 with 1 M NaOH. The 1000 mM HA was added to the TAP medium to make 20 mM pH4.5 HA/TAP. For 5 mM Na^+^ /HEPES and K^+^/HEPES, pH of 10 mM HEPES was adjusted to 7.4 with NaOH or KOH respectively. To make 21 mM Na^+^/EGTA and K^+^/EGTA solutions, 10 mM EGTA was titrated to pH8 with NaOH or KOH. The 5 mM NaCl and KCl solutions were made by dissolving the respective salt in ddw. The 5, 55, and 150 mM Na^+^/HEPES solutions were made by adding NaCl into 5 mM pH7.4 Na^+^/HEPES. The solutions of 30 and 75 mM Ca^2+^/HEPES were made by dissolving CaCl_2_ in 5 mM pH7.4 Na^+^/HEPES.

### Live Cell Imaging and Treatments

EB1-NG in live *Chlamydomonas* cells was imaged with Nikon Eclipse widefield microscope equipped with a short-arc mercury lamp, an FITC-HYQ optical filter set, a CoolSNAP-ES CCD camera and MetaMorph software. Each image was captured as a 16-bit grayscale file with 1 second exposure. Streaming videos were recorded for 100 frames at a rate of 1 frame/second.

Typically cells were resuspended in solutions for 5 min unless indicated otherwise. An aliquot of 5 µl cell suspension was placed on a slide and then covered by an 18 X 18-mm^2^ cover slip. The edges were sealed with nail polish before imaging. For compression experiments, a 3 µl aliquot of cell suspension was placed on a glass slide and then covered with a 22 X 22-mm^2^ cover slip. Cells became gradually compressed by the coverslip as evident by flattened cell body. For pH pulse in a perfusion chamber, an aliquot of 10 µl cells in the TAP medium was placed on a cover slip pre-coated with 5 µl 0.001% poly-L-lysine. The cover slip was then inverted to assemble a perfusion chamber as shown in Figure 2a. The chamber was flushed with 200 µl 20 mM pH4.5 HA/TAP. Subsequently, HA/TAP was replaced by a flush of 200 µl TAP. The entire process was recorded in two consecutive live-stream clips. For this long recording duration, excitation light intensity was reduced to 25% with a neutral density filter. For HA pulse in a diffusion chamber, 40 µl cells in 5 mM pH7.4 Na^+^/HEPES was placed at one side of a diffusion chamber underneath a 40X objective lens (Figure 2a). A live-streaming video was recorded following the injection of 20 µl 100 mM pH2.8 HA through the Vaseline wall to the opposite side of the chamber. For HA bath, a cell pellet from 50 µl liquid culture was resuspended in 50 µl 10 mM pH3 HA. An aliquot of 10 µl cell mixture was placed on a cover slip. The cell-loaded cover slip was inverted to create a perfusion chamber. After a total 5-min exposure to HA, HA was flushed away with an aliquot of 200 µl-indicated fluid and then a video was recorded. To test MT cold lability after recovery from HA bath, a perfusion chamber with treated cells was chilled by ice for 3 minutes. A video was taken immediately afterwards, ~ 20 seconds after the chamber was removed from ice. Each treatment was repeated at least twice in each experiment. Individual experiments were repeated independently 3 times at least.

### Image Analysis

To measure EB1 comet speed, a 40-second substack containing side views of cells were first made by the open source image process software, ImageJ (<u>https://imagej.nih.gov/ij/index.html</u>); and individual comets were analyzed with a Matlab-based particle tracking software, plusTipTracker (Applegate et al. 2011). In each cell that maintained completely quiescent for the tracking period, all tractable comets which transverse at least one third of the cell length were analyzed. The numbers of qualified cells and comets from numerous recordings were indicated. To generate line scans of EB1 intensity at microtubule plus ends, a line tool in ImageJ was used to measure gray values along the length of comets. Relative fluorescence intensity was normalized after calculation by subtracting a background gray value measured next to the comet with the line tool. Histograms were generated with the Microsoft program, Excel. Kymographs were generated with an ImageJ plug-in multiple kymograph (<u>https://www.embl.de/eamnet/html/body_kymograph.html</u>).

### Statistical Analysis

All data are given as mean±SEM (standard error of the mean) and analyzed with Sigmaplot 13.0 (Systat Software, Inc., San Jose, CA). Sample sizes for comet speed measurement are limited by the fact that few cells are entirely quiescent, which is necessary for digital tracking.

## Acknowledgement

This work is supported by Marquette University Startup for P. Yang.

## Supplemental data

Video 1-1 (for Figure 1b) EB1-NG comets in WT cells.

Video 1-2 (for Figure 1f1) Transient bird-cage pattern in WT cells that occurred sporadically during imaging.

Video 1-3 (for Figure 1f2) Disappearance and return of comets in compressed cells following alternate periods of illumination and darkness.

Video 3 (for Figure 3a) WT cells in Na^+^/HEPES after 5-min HA bath.

Figure 4-1 (for Figure 4b) *tub2* cells in HA bath.

Video 4-1 (for Figure 4b) *tub2* cells in Na^+^/HEPES after 5-min HA bath.

Video 4-2 (for Figure 4d). WT cells treated with 5 mM HA.

## References

Akhmanova, A., and Steinmetz, M.O. (2010). Microtubule +TIPs at a glance. J Cell Sci 123, 3415–3419.

Applegate, K.T., Besson, S., Matov, A., Bagonis, M.H., Jaqaman, K., and Danuser, G. (2011). plusTipTracker: Quantitative image analysis software for the measurement of microtubule dynamics. J Struct Biol 176, 168–184.

Arikawa, M., and Suzaki, T. (2002). Reactivation of Ca2+-dependent cytoplasmic contraction in permeabilized cell models of the heliozoon Echinosphaerium akamae. Cell Motil Cytoskeleton 53, 267–272.

Braun, F.J., and Hegemann, P. (1999). Direct measurement of cytosolic calcium and pH in living Chlamydomonas reinhardtii cells. Eur J Cell Biol 78, 199–208.

Buey, R.M., Mohan, R., Leslie, K., Walzthoeni, T., Missimer, J.H., Menzel, A., Bjelic, S., Bargsten, K., Grigoriev, I., Smal, I., Meijering, E., Aebersold, R., Akhmanova, A., and Steinmetz, M.O. (2011). Insights into EB1 structure and the role of its C-terminal domain for discriminating microtubule tips from the lattice. Mol Biol Cell 22, 2912–2923.

Cannon, C., van Adelsberg, J., Kelly, S., and Al-Awqati, Q. (1985). Carbon-dioxide-induced exocytotic insertion of H+ pumps in turtle-bladder luminal membrane: role of cell pH and calcium. Nature 314, 443–446.

Chowdhury, S., Smith, K.W., and Gustin, M.C. (1992). Osmotic stress and the yeast cytoskeleton: phenotype-specific suppression of an actin mutation. J Cell Biol 118, 561–571.

Dhonukshe, P., Laxalt, A.M., Goedhart, J., Gadella, T.W., and Munnik, T. (2003). Phospholipase d activation correlates with microtubule reorganization in living plant cells. Plant Cell 15, 2666–2679.

Dixit, R., and Cyr, R. (2004). Encounters between dynamic cortical microtubules promote ordering of the cortical array through angle-dependent modifications of microtubule behavior. Plant Cell 16, 3274–3284.

Dymek, E.E., Goduti, D., Kramer, T., and Smith, E.F. (2006). A kinesin-like calmodulin-binding protein in Chlamydomonas: evidence for a role in cell division and flagellar functions. J Cell Sci 119, 3107–3116.

Fitzer, S.C., Chung, P., Maccherozzi, F., Dhesi, S.S., Kamenos, N.A., Phoenix, V.R., and Cusack, M. (2016). Biomineral shell formation under ocean acidification: a shift from order to chaos. Sci Rep 6, 21076.

Franck, A.D., Powers, A.F., Gestaut, D.R., Gonen, T., Davis, T.N., and Asbury, C.L. (2007). Tension applied through the Dam1 complex promotes microtubule elongation providing a direct mechanism for length control in mitosis. Nat Cell Biol 9, 832–837.

Fujita, S., Pytela, J., Hotta, T., Kato, T., Hamada, T., Akamatsu, R., Ishida, Y., Kutsuna, N., Hasezawa, S., Nomura, Y., Nakagami, H., and Hashimoto, T. (2013). An atypical tubulin kinase mediates stress-induced microtubule depolymerization in Arabidopsis. Curr Biol 23, 1969–1978.

Fujiu, K., Nakayama, Y., Iida, H., Sokabe, M., and Yoshimura, K. (2011). Mechanoreception in motile flagella of Chlamydomonas. Nat Cell Biol 13, 630–632.

Gagliardi, L.J., and Shain, D.H. (2013). Is intracellular pH a clock for mitosis? Theor Biol Med Model 10, 8.

Gao, X., Zhang, F., Hu, J., Cai, W., Shan, G., Dai, D., Huang, K., and Wang, G. (2016). MicroRNAs modulate adaption to multiple abiotic stresses in Chlamydomonas reinhardtii. Sci Rep 6, 38228.

Gardiner, J. (2013). The evolution and diversification of plant microtubule-associated proteins. Plant J 75, 219–229.

Gardner, M.K., Zanic, M., and Howard, J. (2013). Microtubule catastrophe and rescue. Curr Opin Cell Biol 25, 14–22.

Gibbon, B.C., and Kropf, D.L. (1994). Cytosolic pH Gradients Associated with Tip Growth. Science 263, 1419–1421.

Goldspink, D.A., Gadsby, J.R., Bellett, G., Keynton, J., Tyrrell, B.J., Lund, E.K., Powell, P.P., Thomas, P., and Mogensen, M.M. (2013). The microtubule end-binding protein EB2 is a central regulator of microtubule reorganisation in apico-basal epithelial differentiation. J Cell Sci 126, 4000–4014.

Hardham, A.R. (2013). Microtubules and biotic interactions. Plant J 75, 278–289.

Harris, J.A., Liu, Y., Yang, P., Kner, P., and Lechtreck, K.F. (2016). Single-particle imaging reveals intraflagellar transport-independent transport and accumulation of EB1 in Chlamydomonas flagella. Mol Biol Cell 27, 295–307.

Hashimoto, T. (2015). Microtubules in plants. Arabidopsis Book 13, e0179.

Haswell, E.S., and Verslues, P.E. (2015). The ongoing search for the molecular basis of plant osmosensing. J Gen Physiol 145, 389–394.

Hegemann, P., Berthold, P. (2009). Sensory photoreceptors and light control of flagellar activity. Chapter 13. In: The Chlamydomonas Sourcebook, vol. 3, ed. G.B. Witman: Elsevier, 395–430.

Helenius, J., Brouhard, G., Kalaidzidis, Y., Diez, S., and Howard, J. (2006). The depolymerizing kinesin MCAK uses lattice diffusion to rapidly target microtubule ends. Nature 441, 115–119.

Hepler, P.K. (2016). The Cytoskeleton and Its Regulation by Calcium and Protons. Plant Physiol 170, 3–22.

Hilton, L.K., Meili, F., Buckoll, P.D., Rodriguez-Pike, J.C., Choutka, C.P., Kirschner, J.A., Warner, F., Lethan, M., Garces, F.A., Qi, J., and Quarmby, L.M. (2016). A Forward Genetic Screen and Whole Genome Sequencing Identify Deflagellation Defective Mutants in Chlamydomonas, Including Assignment of ADF1 as a TRP Channel. G3 (Bethesda) 6, 3409–3418.

Hoegh-Guldberg, O., Mumby, P.J., Hooten, A.J., Steneck, R.S., Greenfield, P., Gomez, E., Harvell, C.D., Sale, P.F., Edwards, A.J., Caldeira, K., Knowlton, N., Eakin, C.M., Iglesias-Prieto, R., Muthiga, N., Bradbury, R.H., Dubi, A., and Hatziolos, M.E. (2007). Coral reefs under rapid climate change and ocean acidification. Science 318, 1737–1742.

Horst, C.J., Fishkind, D.J., Pazour, G.J., and Witman, G.B. (1999). An insertional mutant of Chlamydomonas reinhardtii with defective microtubule positioning. Cell Motil Cytoskeleton 44, 143–154.

Husic, H.D., and Tolbert, N.E. (1986). Effect of Osmotic Stress on Carbon Metabolism in Chlamydomonas reinhardtii: Accumulation of Glycerol as an Osmoregulatory Solute. Plant Physiol 82, 594–596.

Karr, T.L., Kristofferson, D., and Purich, D.L. (1980). Calcium ion induces endwise depolymerization of bovine brain microtubules. J Biol Chem 255, 11853–11856.

Kubo, T., Kaida, S., Abe, J., Saito, T., Fukuzawa, H., and Matsuda, Y. (2009). The Chlamydomonas hatching enzyme, sporangin, is expressed in specific phases of the cell cycle and is localized to the flagella of daughter cells within the sporangial cell wall. Plant Cell Physiol 50, 572–583.

Kumar, P., and Wittmann, T. (2012). +TIPs: SxIPping along microtubule ends. Trends Cell Biol 22, 418–428.

Lang, M., Stober, F., and Lichtenthaler, H.K. (1991). Fluorescence emission spectra of plant leaves and plant constituents. Radiat Environ Biophys 30, 333–347.

Lefebvre, P.A., Nordstrom, S.A., Moulder, J.E., and Rosenbaum, J.L. (1978). Flagellar elongation and shortening in Chlamydomonas. IV. Effects of flagellar detachment, regeneration, and resorption on the induction of flagellar protein synthesis. J Cell Biol 78, 8–27.

Liang, X., Madrid, J., and Howard, J. (2014). The microtubule-based cytoskeleton is a component of a mechanical signaling pathway in fly campaniform receptors. Biophys J 107, 2767–2774.

Lieuvin, A., Labbe, J.C., Doree, M., and Job, D. (1994). Intrinsic microtubule stability in interphase cells. J Cell Biol 124, 985–996.

Malhotra, B., and Glass, A. (1995). Potassium Fluxes in Chlamydomonas reinhardtii (II. Compartmental Analysis). Plant Physiol 108, 1537–1545.

Matov, A., Applegate, K., Kumar, P., Thoma, C., Krek, W., Danuser, G., and Wittmann, T. (2010). Analysis of microtubule dynamic instability using a plus-end growth marker. Nat Methods 7, 761–768.

Maurer, S.P., Cade, N.I., Bohner, G., Gustafsson, N., Boutant, E., and Surrey, T. (2014). EB1 accelerates two conformational transitions important for microtubule maturation and dynamics. Curr Biol 24, 372–384.

Maurer, S.P., Fourniol, F.J., Bohner, G., Moores, C.A., and Surrey, T. (2012). EBs recognize a nucleotide-dependent structural cap at growing microtubule ends. Cell 149, 371–382.

Merchant, S.S., Prochnik, S.E., Vallon, O., Harris, E.H., Karpowicz, S.J., Witman, G.B., Terry, A., Salamov, A., Fritz-Laylin, L.K., Marechal-Drouard, L., Marshall, W.F., Qu, L.H., Nelson, D.R., Sanderfoot, A.A., Spalding, M.H., Kapitonov, V.V., Ren, Q., Ferris, P., Lindquist, E., Shapiro, H., Lucas, S.M., Grimwood, J., Schmutz, J., Cardol, P., Cerutti, H., Chanfreau, G., Chen, C.L., Cognat, V., Croft, M.T., Dent, R., Dutcher, S., Fernandez, E., Fukuzawa, H., Gonzalez-Ballester, D., Gonzalez-Halphen, D., Hallmann, A., Hanikenne, M., Hippler, M., Inwood, W., Jabbari, K., Kalanon, M., Kuras, R., Lefebvre, P.A., Lemaire, S.D., Lobanov, A.V., Lohr, M., Manuell, A., Meier, I., Mets, L., Mittag, M., Mittelmeier, T., Moroney, J.V., Moseley, J., Napoli, C., Nedelcu, A.M., Niyogi, K., Novoselov, S.V., Paulsen, I.T., Pazour, G., Purton, S., Ral, J.P., Riano-Pachon, D.M., Riekhof, W., Rymarquis, L., Schroda, M., Stern, D., Umen, J., Willows, R., Wilson, N., Zimmer, S.L., Allmer, J., Balk, J., Bisova, K., Chen, C.J., Elias, M., Gendler, K., Hauser, C., Lamb, M.R., Ledford, H., Long, J.C., Minagawa, J., Page, M.D., Pan, J., Pootakham, W., Roje, S., Rose, A., Stahlberg, E., Terauchi, A.M., Yang, P., Ball, S., Bowler, C., Dieckmann, C.L., Gladyshev, V.N., Green, P., Jorgensen, R., Mayfield, S., Mueller-Roeber, B., Rajamani, S., Sayre, R.T., Brokstein, P., Dubchak, I., Goodstein, D., Hornick, L., Huang, Y.W., Jhaveri, J., Luo, Y., Martinez, D., Ngau, W.C., Otillar, B., Poliakov, A., Porter, A., Szajkowski, L., Werner, G., Zhou, K., Grigoriev, I.V., Rokhsar, D.S., and Grossman, A.R. (2007). The Chlamydomonas genome reveals the evolution of key animal and plant functions. Science 318, 245–250.

Miller, D.J. (1979). Are cardiac muscle cells 'skinned' by EGTA or EDTA? Nature 277, 142–143.

Mitchison, T., and Kirschner, M. (1984). Dynamic instability of microtubule growth. Nature 312, 237–242.

Mittelmeier, T.M., Boyd, J.S., Lamb, M.R., and Dieckmann, C.L. (2011). Asymmetric properties of the Chlamydomonas reinhardtii cytoskeleton direct rhodopsin photoreceptor localization. J Cell Biol 193, 741–753.

Muller-Reichert, T., Chretien, D., Severin, F., and Hyman, A.A. (1998). Structural changes at microtubule ends accompanying GTP hydrolysis: information from a slowly hydrolyzable analogue of GTP, guanylyl (alpha,beta)methylenediphosphonate. Proc Natl Acad Sci U S A 95, 3661–3666.

Nagel, G., Ollig, D., Fuhrmann, M., Kateriya, S., Musti, A.M., Bamberg, E., and Hegemann, P. (2002). Channelrhodopsin-1: a light-gated proton channel in green algae. Science 296, 2395–2398.

Nagel, G., Szellas, T., Huhn, W., Kateriya, S., Adeishvili, N., Berthold, P., Ollig, D., Hegemann, P., and Bamberg, E. (2003). Channelrhodopsin-2, a directly light-gated cation-selective membrane channel. Proc Natl Acad Sci U S A 100, 13940–13945.

O'Brien, E.T., Salmon, E.D., and Erickson, H.P. (1997). How calcium causes microtubule depolymerization. Cell Motil Cytoskeleton 36, 125–135.

Oda, Y. (2015). Cortical microtubule rearrangements and cell wall patterning. Front Plant Sci 6, 236.

Olmsted, J.B., and Borisy, G.G. (1975). Ionic and nucleotide requirements for microtubule polymerization in vitro. Biochemistry 14, 2996–3005.

Paredez, A.R., Somerville, C.R., and Ehrhardt, D.W. (2006). Visualization of cellulose synthase demonstrates functional association with microtubules. Science 312, 1491–1495.

Pedersen, L.B., Geimer, S., Sloboda, R.D., and Rosenbaum, J.L. (2003). The Microtubule plus end-tracking protein EB1 is localized to the flagellar tip and basal bodies in Chlamydomonas reinhardtii. Curr Biol 13, 1969–1974.

Perrineau, M.M., Zelzion, E., Gross, J., Price, D.C., Boyd, J., and Bhattacharya, D. (2014). Evolution of salt tolerance in a laboratory reared population of Chlamydomonas reinhardtii. Environ Microbiol 16, 1755–1766.

Picariello, T., Valentine, M.S., Yano, J., and Van Houten, J. (2014). Reduction of meckelin leads to general loss of cilia, ciliary microtubule misalignment and distorted cell surface organization. Cilia 3, 2.

Piehl, M., Tulu, U.S., Wadsworth, P., and Cassimeris, L. (2004). Centrosome maturation: measurement of microtubule nucleation throughout the cell cycle by using GFP-tagged EB1. Proc Natl Acad Sci U S A 101, 1584–1588.

Pittman, J.K., Edmond, C., Sunderland, P.A., and Bray, C.M. (2009). A cation-regulated and proton gradient-dependent cation transporter from Chlamydomonas reinhardtii has a role in calcium and sodium homeostasis. J Biol Chem 284, 525–533.

Pohl, H.R., Wheeler, J.S., and Murray, H.E. (2013). Sodium and potassium in health and disease. Met Ions Life Sci 13, 29–47.

Prachayasittikul, V., Isarankura-Na-Ayudhya, C., Tantimongcolwat, T., Nantasenamat, C., and Galla, H.J. (2007). EDTA-induced membrane fluidization and destabilization: biophysical studies on artificial lipid membranes. Acta Biochim Biophys Sin (Shanghai) 39, 901–913.

Prager-Khoutorsky, M., Khoutorsky, A., and Bourque, C.W. (2014). Unique interweaved microtubule scaffold mediates osmosensory transduction via physical interaction with TRPV1. Neuron 83, 866–878.

Quarmby, L.M. (1996). Ca2+ influx activated by low pH in Chlamydomonas. J Gen Physiol 108, 351–361.

Quarmby, L.M. (2009). Deflagellation. Chapter 3. In: The Chlamydomonas Sourcebook, vol. 3, ed. G.B. Witman: Elsevier, 43–69.

Rasala, B.A., Barrera, D.J., Ng, J., Plucinak, T.M., Rosenberg, J.N., Weeks, D.P., Oyler, G.A., Peterson, T.C., Haerizadeh, F., and Mayfield, S.P. (2013). Expanding the spectral palette of fluorescent proteins for the green microalga Chlamydomonas reinhardtii. Plant J 74, 545–556.

Raven, J., Caldeira, K., et al. (2005). Ocean acidification due to increasing atmospheric carbon dioxide. The Royal Society.

Robertson, A.M., and Hagan, I.M. (2008). Stress-regulated kinase pathways in the recovery of tip growth and microtubule dynamics following osmotic stress in S. pombe. J Cell Sci 121, 4055–4068.

Rogers, S.L., Rogers, G.C., Sharp, D.J., and Vale, R.D. (2002). Drosophila EB1 is important for proper assembly, dynamics, and positioning of the mitotic spindle. J Cell Biol 158, 873–884.

Ronkin, R.R., Buretz, K.M. (1962). Sodium and potassium in normal and paralyzed Chlamydomonas. Journal of protozoology 7, 109–114.

Rovini, A., Gauthier, G., Berges, R., Kruczynski, A., Braguer, D., and Honore, S. (2013). Anti-migratory effect of vinflunine in endothelial and glioblastoma cells is associated with changes in EB1 C-terminal detyrosinated/tyrosinated status. PLoS One 8, e65694.

Ruffin, V.A., Salameh, A.I., Boron, W.F., and Parker, M.D. (2014). Intracellular pH regulation by acid-base transporters in mammalian neurons. Front Physiol 5, 43.

Sayas, C.L., and Avila, J. (2014). Crosstalk between axonal classical microtubule-associated proteins and end binding proteins during axon extension: possible implications in neurodegeneration. J Alzheimers Dis 40 Suppl 1, S17–22.

Schatten, G., Bestor, T., Balczon, R., Henson, J., and Schatten, H. (1985). Intracellular pH shift leads to microtubule assembly and microtubule-mediated motility during sea urchin fertilization: correlations between elevated intracellular pH and microtubule activity and depressed intracellular pH and microtubule disassembly. Eur J Cell Biol 36, 116–127.

Schibler, M.J., and Huang, B. (1991). The colR4 and colR15 beta-tubulin mutations in Chlamydomonas reinhardtii confer altered sensitivities to microtubule inhibitors and herbicides by enhancing microtubule stability. J Cell Biol 113, 605–614.

Serrano, L., Valencia, A., Caballero, R., and Avila, J. (1986). Localization of the high affinity calcium-binding site on tubulin molecule. J Biol Chem 261, 7076–7081.

Shaner, N.C., Lambert, G.G., Chammas, A., Ni, Y., Cranfill, P.J., Baird, M.A., Sell, B.R., Allen, J.R., Day, R.N., Israelsson, M., Davidson, M.W., and Wang, J. (2013). A bright monomeric green fluorescent protein derived from Branchiostoma lanceolatum. Nat Methods 10, 407–409.

Shoji, T., Suzuki, K., Abe, T., Kaneko, Y., Shi, H., Zhu, J.K., Rus, A., Hasegawa, P.M., and Hashimoto, T. (2006). Salt stress affects cortical microtubule organization and helical growth in Arabidopsis. Plant Cell Physiol 47, 1158–1168.

Solomon, F. (1977). Binding sites for calcium on tubulin. Biochemistry 16, 358–363.

Song, Y., and Brady, S.T. (2014). Post-translational modifications of tubulin: pathways to functional diversity of microtubules. Trends Cell Biol 25, 125–136.

Su, L.K., Burrell, M., Hill, D.E., Gyuris, J., Brent, R., Wiltshire, R., Trent, J., Vogelstein, B., and Kinzler, K.W. (1995). APC binds to the novel protein EB1. Cancer Res 55, 2972–2977.

Takouridis, S.J., Tribe, D.E., Gras, S.L., and Martin, G.J. (2015). The selective breeding of the freshwater microalga Chlamydomonas reinhardtii for growth in salinity. Bioresour Technol 184, 18–22.

Taylor, A.R., Brownlee, C., and Wheeler, G.L. (2012). Proton channels in algae: reasons to be excited. Trends Plant Sci 17, 675–684.

Tazawa, M., and Shimmen, T. (1983). Cell motility and ionic relations in Characean cells as revealed by internal perfusion and cell models. Int Rev Cytology 21, 1535–1540.

Tirnauer, J.S., Grego, S., Salmon, E.D., and Mitchison, T.J. (2002). EB1-microtubule interactions in Xenopus egg extracts: role of EB1 in microtubule stabilization and mechanisms of targeting to microtubules. Mol Biol Cell 13, 3614–3626.

Tortosa, E., Galjart, N., Avila, J., and Sayas, C.L. (2013). MAP1B regulates microtubule dynamics by sequestering EB1/3 in the cytosol of developing neuronal cells. Embo J 32, 1293–1306.

Van Damme, D., Bouget, F.Y., Van Poucke, K., Inze, D., and Geelen, D. (2004). Molecular dissection of plant cytokinesis and phragmoplast structure: a survey of GFP-tagged proteins. Plant J 40, 386–398.

Vitre, B., Coquelle, F.M., Heichette, C., Garnier, C., Chretien, D., and Arnal, I. (2008). EB1 regulates microtubule dynamics and tubulin sheet closure in vitro. Nat Cell Biol 10, 415–421.

Waldbusser, G.G., Bruner, E.L., Haley, B.A., Hales, B., Langdon, C.J., Prahl, F.G. (2013). A developmental and energetic basis linking larval oyster shell formation to acidification sensitivity Geophysical Research Letters 40, 2171–2176.

Wang, C., Li, J., and Yuan, M. (2007). Salt tolerance requires cortical microtubule reorganization in Arabidopsis. Plant Cell Physiol 48, 1534–1547.

Wang, C., Zhang, L., and Chen, W. (2011). Plant cortical microtubules are putative sensors under abiotic stresses. Biochemistry (Mosc) 76, 320–326.

Wang, L., Piao, T., Cao, M., Qin, T., Huang, L., Deng, H., Mao, T., and Pan, J. (2013). Flagellar regeneration requires cytoplasmic microtubule depolymerization and kinesin-13. J Cell Sci 126, 1531–1540.

Wang, S., Kurepa, J., Hashimoto, T., and Smalle, J.A. (2011). Salt stress-induced disassembly of Arabidopsis cortical microtubule arrays involves 26S proteasome-dependent degradation of SPIRAL1. Plant Cell 23, 3412–3427.

Weaver, B.A. (2014). How Taxol/paclitaxel kills cancer cells. Mol Biol Cell 25, 2677–2681.

Weisenberg, R.C. (1972). Microtubule formation in vitro in solutions containing low calcium concentrations. Science 177, 1104–1105.

Wheeler, G.L., Joint, I., and Brownlee, C. (2008). Rapid spatiotemporal patterning of cytosolic Ca2+ underlies flagellar excision in Chlamydomonas reinhardtii. Plant J 53, 401–413.

Wolff, J., Sackett, D.L., and Knipling, L. (1996). Cation selective promotion of tubulin polymerization by alkali metal chlorides. Protein Sci 5, 2020–2028.

Yan, X., Habedanck, R., and Nigg, E.A. (2006). A complex of two centrosomal proteins, CAP350 and FOP, cooperates with EB1 in microtubule anchoring. Mol Biol Cell 17, 634–644.

Zanic, M., Stear, J.H., Hyman, A.A., and Howard, J. (2009). EB1 recognizes the nucleotide state of tubulin in the microtubule lattice. PLoS One 4, e7585.

Zhang, Q., Lin, F., Mao, T., Nie, J., Yan, M., Yuan, M., and Zhang, W. (2012). Phosphatidic acid regulates microtubule organization by interacting with MAP65-1 in response to salt stress in Arabidopsis. Plant Cell 24, 4555–4576.

